# Distinguishing cophylogenetic signal from phylogenetic congruence clarifies the interplay between evolutionary history and species interactions

**DOI:** 10.1101/2023.07.21.550001

**Authors:** Benoît Perez-Lamarque, Hélène Morlon

## Abstract

Interspecific interactions, including host-symbiont associations, can profoundly affect the evolution of the interacting species. Given the phylogenies of host and symbiont clades and knowledge of which host species interact with which symbiont, two questions are often asked: “Do closely related hosts interact with closely related symbionts?” and “Do host and symbiont phylogenies mirror one another?”. These questions are intertwined and can even collapse under specific situations, such that they are often confused one with the other. However, in most situations, a positive answer to the first question, hereafter referred to as “cophylogenetic signal”, does not imply a close match between the host and symbiont phylogenies. It suggests only that past evolutionary history has contributed to shaping present-day interactions, which can arise, for example, through present-day trait matching, or from a single ancient vicariance event that increases the probability that closely related species overlap geographically. A positive answer to the second, referred to as “phylogenetic congruence”, is more restrictive as it suggests a close match between the two phylogenies, which may happen, for example, if symbiont diversification tracks host diversification or if the diversifications of the two clades were subject to the same succession of vicariance events.

Here we apply a set of methods (ParaFit, PACo, and eMPRess), which significance is often interpreted as evidence for phylogenetic congruence, to simulations under three biologically realistic scenarios of trait matching, a single ancient vicariance event, and phylogenetic tracking with frequent cospeciation events. The latter is the only scenario that generates phylogenetic congruence, whereas the first two generate a cophylogenetic signal in the absence of phylogenetic congruence. We find that tests of global-fit methods (ParaFit and PACo) are significant under the three scenarios, whereas tests of event-based methods (eMPRess) are only significant under the scenario of phylogenetic tracking. Therefore, significant results from global-fit methods should be interpreted in terms of cophylogenetic signal and not phylogenetic congruence; such significant results can arise under scenarios when hosts and symbionts had independent evolutionary histories. Conversely, significant results from event-based methods suggest a strong form of dependency between hosts and symbionts evolutionary histories. Clarifying the patterns detected by different cophylogenetic methods is key to understanding how interspecific interactions shape and are shaped by evolution.

## Introduction

Antagonistic or mutualistic interactions, such as parasitism, herbivory, seed dispersal, or pollination, are key components of ecological communities (Bascompte and Jordano 2013; Mittelbach and McGill 2019). Patterns of interactions (*i.e.* who interacts with whom) are shaped by evolutionary history through a variety of processes. For example, interspecific interactions may be constrained to species with matching traits, as frequently observed in pollination or host-parasite interaction networks (Muchhala and Thomson 2009; Morand et al. 2015), in which case species evolutionary history matters as soon as the traits involved in the interactions are evolutionarily conserved. Interspecific interactions may also be constrained by historical contingencies such as past dispersal and/or geographic events: for instance, if lineages have been geographically isolated following a vicariance event, these lineages do not interact simply because they do not co-occur (Althoff et al. 2014; Perez-Lamarque et al. 2022a). Interspecific interactions can also be transmitted from generation to generation on evolutionary time scales, as in the case of symbionts that are vertically transmitted from parental to descendant hosts (Bright and Bulgheresi 2010). Reciprocally, interactions affect the evolution of the interacting species, for instance through disruptive selection, stabilizing selection, or coevolution. Over macroevolutionary scales, such effects can leave an imprint on the phylogenetic trees of the interacting clades (Harmon et al. 2019; Hembry and Weber 2020; Hayward et al. 2021). An extreme case corresponds to phylogenetic tracking, which can happen, for example, when host speciation events lead to the subsequent speciation of symbionts that are closely associated with them (Fahrenholz 1912). Analyzing interacting species through the lens of their past evolutionary history is therefore fundamental for understanding how interspecific interactions shape and are shaped by evolution.

Given host-symbiont cophylogenetic data, *i.e.* a phylogenetic tree for both the host and the symbiont clades and knowledge of which host species interact with which symbiont, two patterns are often investigated (Supplementary Box 1): (i) whether closely-related hosts interact with closely-related symbionts, hereafter referred to as “cophylogenetic signal”, and (ii) whether host and symbiont phylogenies mirror one another (with pairs of interacting hosts and symbionts that tend to occupy similar positions in the two trees), hereafter referred to as “phylogenetic congruence” (Blasco-Costa et al. 2021). By definition, phylogenetic congruence can occur only when the number of host and symbiont species is similar, with mostly ‘one-to-one’ interactions between one host and one symbiont species. In this case, patterns of cophylogenetic signal and phylogenetic congruence tend to collapse, such that they can be studied interchangeably. In this context, some cophylogenetic methods, referred to as global-fit methods, have been developed to test for phylogenetic congruence using linear algebra techniques, such as the fourth-corner statistics (ParaFit, Legendre et al. 2002) and procrustean superposition (PACo, Balbuena et al. 2013). Numerous cophylogenetic systems observed in nature, however, are characterized by ‘many-to-many’ interactions, as illustrated by densely-connected networks of interspecific interactions (Ronquist and Nylin 1990; Bascompte and Jordano 2013; Pichon et al. 2023). The same global fit methods have been regularly applied to such systems, and interpreted as tests of phylogenetic congruence (see *e.g.* Fuzessy et al. 2022; Suzuki et al. 2022 for recent examples). If, under these situations, global-fit methods actually do not measure phylogenetic congruence, this can lead to a misinterpretation of the results.

Patterns of cophylogenetic signal and phylogenetic congruence can reflect different processes (Fig. 1). The occurrence of cophylogenetic signal simply suggests that past evolutionary history has contributed to shaping present-day interactions. For instance, a cophylogenetic signal can arise if interactions are shaped by evolutionarily conserved traits (Fig. 1a “trait matching”) or by past biogeographic events (Fig. 1b “vicariance”), even if the diversification of the host and symbiont clades was not influenced by the interactions between host and symbiont species. The occurrence of phylogenetic congruence, on the other hand, suggests a potential (co-)dependency between the host and symbiont diversifications, causing both phylogenetic trees to look alike. For instance, phylogenetic congruence can emerge (i) if the diversification of vertically transmitted symbionts tracks host diversification, generating a pattern of concomitant diversification events happening in both host and symbiont clades, referred to as “codiversification” (Fig. 1c “phylogenetic tracking”), (ii) if the diversifications of the two clades were subject to the same succession of vicariance events, also resulting in a pattern of concomitant codiversification, or (iii) under preferential host switching, *i.e.* if symbionts diversify by preferentially transferring to closely-related host species, generating a pattern of phylogenetic congruence without concomitant divergence times, referred to as “pseudo-codiversification” (de Vienne et al. 2013; Althoff et al. 2014). Coevolution, *i.e.* the reciprocal evolutionary changes in interacting lineages induced by selective pressures exerted by one another, may also generate phylogenetic congruence in some cases (*e.g.* aphid-associated bacterial endosymbionts (Jousselin et al. 2009)), but is not a necessary nor sufficient condition for observing phylogenetic congruence (Poisot 2015). Phylogenetic congruence implies cophylogenetic signal, but the reverse is not true.

**Figure 1:**
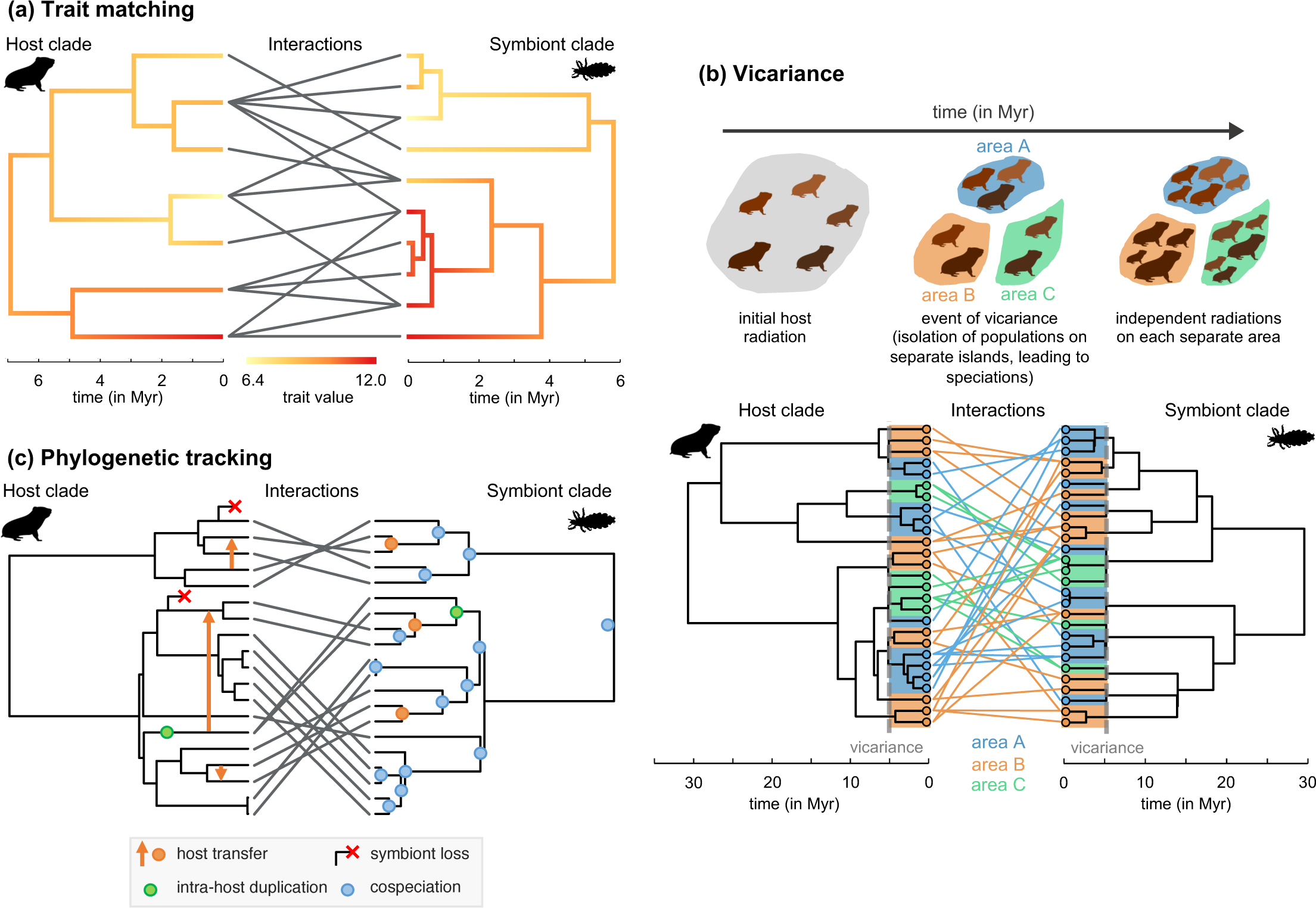
Three mock examples of host-symbiont systems (here represented as gophers and lice) that generate a cophylogenetic signal. Only the third example also generates phylogenetic congruence. **(a) Trait matching.** Here host and symbiont phylogenetic trees evolve independently. Traits also evolve independently on each phylogeny following Brownian motion processes, which results in a phylogenetic signal in species traits (*i.e.* closely related species tend to have similar trait values; see the color gradient). Host-symbiont interactions at present are more likely between species having complementary traits, following a trait-matching expression. Because of trait matching, closely related host species interact with closely related symbiont species (*i.e.* cophylogenetic signal), although the phylogenetic trees are independent (*i.e.* no phylogenetic congruence). **(b) Vicariance:** Here hosts and symbionts interact at random as long as they occupy the same biogeographic area. They occupy a single area until a vicariance event (*i.e.* the formation of a biogeographic barrier) splits this area into three separate areas, isolating populations and leading to speciation (a species occupying two areas at the time of vicariance immediately experiences allopatric speciation). Following the vicariance event, each host and symbiont lineage radiates independently on its area, without dispersal between areas, which results in a phylogenetic signal in biogeographic repartition (*i.e.* closely related species tend live on the same area). Because interactions happen at random within each area, this scenario generates cophylogenetic signal without phylogenetic congruence. **(c) Phylogenetic tracking:** Here symbiont species are vertically transmitted over long-time scales along host lineages and hosts speciations concomitantly lead to symbiont speciations (“cospeciation”) resulting in a pattern of codiversification. In addition, symbiont lineages can experience horizontal host transfers from a donor host to a receiver host (with replacement of the previous symbiont lineage), intra-host duplication, and host lineages can lose their symbionts. This scenario generates both cophylogenetic signal and phylogenetic congruence. The greater the number of host transfers, intra-host duplications, and symbiont loss, the lower the phylogenetic congruence, as these events disrupt the symbiont phylogeny with regard to the host phylogeny (Ronquist 2003a,b).

Here, in an effort to clarify the conclusions that can be drawn from various cophylogenetic methods, we analyze their outputs under various evolutionary scenarios, using simulations. Cophylogenetic methods can be divided into two main types of approaches (see de Vienne et al. (2013) and Dismukes et al. (2022) for reviews of these methods): the global-fit methods mentioned above, such as ParaFit and PACo, and the event-based methods, such as TreeMap (Page 1994a, 1995), TreeFitter (Ronquist 2003a), Jane (Conow et al. 2010), or eMPRess (Santichaivekin et al. 2021). Event-based methods try to reconciliate the host and symbiont phylogenies by fitting a set of reconciliation events (e.g. cospeciation, host transfer, intra-host duplication, or symbiont loss) to the symbiont phylogeny (Page 1994b; Ronquist 2003b). We have shown before, in a different context, that global-fit and event-based methods can sometimes output contrasting results (Perez-Lamarque and Morlon 2023), suggesting that they measure different patterns. We evaluate here their outputs under three biologically realistic scenarios of host-symbiont evolution by trait matching, vicariance, and phylogenetic tracking. The latter is the only scenario that generates phylogenetic congruence when simulating frequent cospeciations, whereas the first two generate a cophylogenetic signal in the absence of phylogenetic congruence. We find that global-fit methods provide a test of cophylogenetic signal rather than phylogenetic congruence, whereas event-based methods provide a test of phylogenetic congruence, and discuss the implications of these results for the study of cophylogenetic systems.

### Simulations of three biologically realistic scenarios

In the first simulated scenario, we assumed that present-day host-symbiont interactions are more likely between species having complementary traits following a trait-matching expression with a unidimensional continuous trait (Fig. 1a – “trait matching”; Supplementary Methods 1). We independently simulated two phylogenetic trees for the host and symbiont clades using a birth-death model and on each tree, we then independently simulated the evolution of traits modulating present-day interactions. By using a Brownian motion for trait evolution, closely-related hosts and closely-related symbionts tend to have similar trait values (*i.e.* phylogenetic signal in species traits). Finally, we assumed that the degree of specialization of the symbionts (*i.e.* the number of hosts that a given symbiont interacts with) follows a Poisson distribution with parameter λ =1.5 and attributed the present-day host-symbiont interacting pairs following a trait-matching expression. As a result, closely related host species interact with closely related symbiont species (*i.e.* cophylogenetic signal), although the phylogenetic trees were simulated independently and are therefore not congruent. We generated 1,000 simulations with varying clade sizes (from 10 to 200 species per clade; Supplementary Methods 1) and replicated the simulations with fewer host species associated with each symbiont species by using a Poisson distribution with parameter λ=1.

In the second simulated scenario, we assumed that host and symbiont species interact at random as long as they occupy the same biogeographic area. At first, all hosts and symbionts simultaneously occupy a single area and diversify independently, until a vicariance event splits the area into three separate areas (Fig. 1b – “vicariance”; Supplementary Methods 1). Half of the host and symbiont species are then isolated in separate areas; the other half experiences allopatric speciation as their population is split into two. Following the vicariance event, each host and symbiont lineage diversifies independently in its area, resulting in a phylogenetic signal in biogeographic repartition (*i.e.* closely related species tend to occupy the same area). Although the host and symbiont diversifications are not independent (they both undergo a burst of speciation events at the time of vicariance), the host and symbiont phylogenetic trees are not congruent as they experienced different events of diversification before and after the vicariance (Fig. 1b). Finally, we assumed that the degree of specialization of the symbionts follows a Poisson distribution with parameter λ=1.5 and randomly attributed host-symbiont present-day interactions within each area. This scenario thus produces cophylogenetic signal but no phylogenetic congruence. We generated 1,000 simulations with varying clade sizes (Supplementary Methods 1) and replicated the simulations with fewer host species associated with each symbiont species (λ=1).

In the third simulated scenario, we assumed that the symbiont diversification tracks the host diversification. Symbionts species are vertically transmitted over long-time scales from host generation to host generation (at the level of host individual or at the level of the whole host lineage) and cospeciate at host speciation events resulting in codiversification (Fig. 1c – “phylogenetic tracking”; Supplementary Methods 1). In addition, we assumed that symbiont lineages experience a given number of host transfers from a donor host to a receiver host (with replacement of the previous symbiont lineage); this number was uniformly sampled between 0 and half the number of extant host species. Finally, intra-host duplication occurs at rate 0.001 event per million year per lineage and host lineages can lose their symbionts with a probability of 0.1 at present. The latter processes (transfer, duplication and loss) dampen but maintain the phylogenetic congruence between host and symbiont phylogenies, rendering it more realistic. This third scenario of phylogenetic tracking with frequent cospeciations produces both cophylogenetic signal and phylogenetic congruence. We generated 1,000 simulations with varying clade sizes (from 10 to 200 host species; Supplementary Methods 1). We also replicated the simulations with less cospeciations and more host transfers (number uniformly sampled between 50% and 75% of the number of extant host species) and intra-host duplications (rate of 0.0015 event per million year per lineage). Under these simulations of phylogenetic tracking with infrequent cospeciations, the pattern of phylogenetic congruence is greatly erased as cospeciation events represent a minority of simulated events.

We therefore obtained a total of 6,000 simulations with different host and symbiont species richness values and ratios of one-to-one interactions (Supplementary Figs. S1 & S2). As expected, our simulations under phylogenetic tracking produced systems with a much higher proportion of one-to-one interactions than simulations under trait matching and vicariance, which were mainly constituted of many-to-many interactions, especially under low symbiont specialization ( λ=1.5, Supplementary Fig. S2).

### Fitting cophylogenetic methods

For each simulated cophylogenetic data, we first applied the global-fit methods ParaFit and PACo using the functions *parafit* and *PACo* from the R-packages ape (Paradis et al. 2004) and paco (Hutchinson et al. 2017) respectively, amended to avoid technical issues when the number of host or symbiont species is low (Perez-Lamarque and Morlon 2023). We measured the strength of the cophylogenetic signal using the “ParaFit global statistic” in ParaFit, whereas, in PACo, we ran the symmetric option for the procrustean superposition to obtain R^2^, defined as R^2^ = 1 – m^2^, where m^2^ is the sum of the squared residuals of the symmetric procrustean superposition (Blasco-Costa et al. 2021). R^2^ is comprised between 0 and 1, and R^2^ close to 0 indicates low cophylogenetic signal, whereas R^2^ close to 1 indicates high cophylogenetic signal. Following Legendre et al. (2002) and Hutchinson et al. (2017), the significance of the ParaFit and PACo tests was first evaluated using 10,000 randomizations obtained by independently shuffling which host species are associated with each symbiont species, hereafter referred to as “null model 1”. Second, following Ronquist (1998) and Sanmartín and Ronquist (2004), we assessed their significance by randomly shuffling the host species labels, hereafter referred to as “null model 2”. We also investigated whether the effect size of the global-fit methods, *i.e.* values of the ParaFit global statistic and PACo’s R^2^, can be used as indicators of phylogenetic congruence (Blasco-Costa et al. 2021). Finally, we tested whether global-fit approaches were more likely to be significant when the ratio of one-to-one interactions was low using generalized linear models.

Second, we applied the event-based method eMPRess to each simulation (Santichaivekin et al. 2021). eMPRess reconciles the host and symbiont tree topologies by using maximum parsimony to fit events of host transfers, duplications, and losses; each event being associated with a given cost. We chose eMPRess over all the other existing event-based methods, as it can be automatically and rapidly run using the command line. As with most event-based methods, eMPRess does not handle symbiont species that interact with multiple hosts (Dismukes et al. 2022). However, symbionts often interact with several hosts in nature, and also in our simulated scenarios of trait matching and vicariance. We thus tested two strategies for running eMPRess following Sanmartín and Ronquist (2002): (i) subsampling one host at random per symbiont species (Su et al. 2022) or (ii) randomly generating bifurcating sub-trees for symbionts with multiple hosts, such that each tip in the symbiont tree is associated with a single host (Satler et al. 2019). We used the command-line version of eMPRess with the commands “python empress_cli.py reconcile” to reconcile the trees and “python empress_cli.py p-value” to assess the significance. We tested 5 different combinations of cost values for duplication (d), host transfer (t) or loss (l) events: (i) d=1, t=1, l=1; (ii) d=4, t=1, l=1; (iii) d=2, t=1, l=2; (iv) d=4, t=2, l=1; or (v) d=2, t=3, l=1. A reconciliation is considered significant if its total cost is lower than 95% of the costs of 1,000 reconciliations obtained after randomly shuffling the host species labels (“null model 2”; Ronquist 1998; Sanmartín and Ronquist 2004). Phylogenetic trees are considered congruent if the reconciliation estimates less host transfer than cospeciation events (Groussin et al. 2017; Perez-Lamarque and Morlon 2023).

### Outputs of cophylogenetic methods

For ParaFit, we found significant tests in 34% of the trait-matching simulations, 45% of the vicariance simulations, and 98% of the phylogenetic tracking (obtained with “null model 1” and λ=1 or 1.5; Table 1). Significant tests for PACo were even more frequent, reaching 48%, 63%, and 99% of the simulations in the three scenarios, respectively (Table 1). Qualitatively similar results were obtained when evaluating the significance by shuffling the host species labels (“null model 2”; Supplementary Table S1). As expected, global-fit tests were more often significant in simulations with higher numbers of host and symbiont species (Supplementary Table S2, Supplementary Figs. S3 & S4). In trait matching and vicariance simulations, when increasing the ratio of one-to-one host-symbiont interactions (*i.e.* reducing the mean number of associated hosts per symbiont by using λ=1), global-fit tests were less often significant (Table 1; Supplementary Table S3; Supplementary Figs. S3 & S4; generalized linear models: p-values<0.05). In other words, global-fit tests are actually more likely to be significant when there are frequent many-to-many host-symbiont interactions, while we would expect the contrary if these tests measured phylogenetic congruence. Overall, these results indicate that global-fit methods tend to measure cophylogenetic signal in general rather than specifically phylogenetic congruence.

In terms of interpretation of the effect size of the global-fit methods, the ParaFit global statistic varies with the total number of species, and is thus difficult to interpret in itself: it cannot be used to distinguish phylogenetic congruence from cophylogenetic signal (Fig. 2; Supplementary Fig. S3). In contrast, for PACo, we found that R^2^ is generally higher than 0.25 when there is a pattern of phylogenetic congruence (in 99% of the scenario of phylogenetic tracking with frequent cospeciations), whereas it is lower than 0.5 when there is a pattern of cophylogenetic signal alone (in 99% of the scenarios of trait-matching or vicariance; Figs. 2 & 3). Hence, a significant test with R^2^ > 0.50 is strong support for phylogenetic congruence (although not definite evidence, it may be possible to obtain higher R^2^ values under scenarios without congruence with different simulation choices), whereas a significant test with R^2^ < 0.25 suggests that the system presents a cophylogenetic signal without phylogenetic congruence (Fig. 3). A significant test with R^2^ values between 0.25 and 0.50 is harder to interpret, as R^2^ tends to correlate positively with the ratio of one-to-one interactions and R^2^ values above 0.25 are sometimes reported in simulations without phylogenetic congruence (Supplementary Fig. S4). PACo alone is therefore often not sufficient to identify a pattern of phylogenetic congruence.

For eMPRess, when subsampling one host per symbiont species, the reconciliation was significant in only 5% of the trait-matching simulations and 16% of the vicariance simulations (the two scenarios generating cophylogenetic signal without phylogenetic congruence), and congruent in none (Table 1). When generating random bifurcations in the symbiont tree, it was significant in 34% and 80% of the trait-matching and vicariance simulations, respectively, but congruent in none (Table 1).

Indeed, eMPRess estimated on average 5 times more host transfers than cospeciations for both scenarios of trait matching and vicariance (Table 1; Fig. 2; Supplementary Fig. S5). In contrast, when simulating phylogenetic tracking with a majority of cospeciation events, eMPRess gave significant reconciliations in 100% of the simulations, 92% of which were congruent (Table 1). Results were qualitatively similar when choosing alternative event costs (Supplementary Tables S4, S5, & S6). In all simulated scenarios, significant reconciliations were more frequent when the number of host and symbiont species were larger (Supplementary Tables S4, S5, & S6). When simulating phylogenetic tracking with infrequent cospeciations, eMPRess reconciliations were still significant in 99% of the simulations, but congruent in only 34% of them; this is expected, given that the number of cospeciations is lower than the number of host transfers in these simulations (Fig. 2; Table 1; Supplementary Tables S7). Overall, our findings suggest that event-based methods can specifically detect patterns of phylogenetic congruence and distinguish them from a “simple” cophylogenetic signal.

**Table 1:** ParaFit and PACo (global-fit methods) are often significant in all simulated scenarios, including scenarios of trait matching and vicariance that do not generate phylogenetic congruence, whereas eMPRess (an event-based method) supports phylogenetic congruence only under scenarios of phylogenetic tracking with frequent cospeciations, which generate phylogenetic congruence: This table indicates the percentages of simulations for which ParaFit, PACo, or eMPRess output a significant test, under three scenarios (Fig. 1): (a) present-day interactions dictated by trait matching (with more (ϕ\=1.5) or less (ϕ\=1) host species associated with each symbiont species), (b) present-day interactions at random following a single vicariance event (with more (ϕ\=1.5) or less (ϕ\=1) host species associated with each symbiont species), or (c) present-day interactions resulting from phylogenetic tracking (with frequent or infrequent cospeciation events). For eMPRess, we report the percentage of significant reconciliations based on permutations alone (P) or based on selecting among these the ones that have more cospeciation than host transfer events (P+C). We consider eMPRess to support phylogenetic congruence when conditions P and C are met (in bold). Results by clade size are given in Supplementary Tables S1-S7. eMPRess results correspond to host-symbiont reconciliations ran with relative costs d=4, t=1, and l=1 for duplications, host transfers, and losses, respectively (other costs combinations provided qualitatively similar results – see Supplementary Tables S4-S6).

| Methods |  | (a)<br>Trait matching |  | (b)<br>Vicariance |  | (c)<br>Phylogenetic tracking |  |
| --- | --- | --- | --- | --- | --- | --- | --- |
| | | $\lambda=1.5$ | $\lambda=1$ | $\lambda=1.5$ | $\lambda=1$ | Frequent co-speciations | Infrequent co-speciations |
| Global-fit methods | Percentage of significant tests using ParaFit | 39% | 29% | 52% | 38% | 99% | 96% |
|  | Percentage of significant tests using PACo | 50% | 45% | 68% | 59% | 100% | 99% |
| Event-based methods | Percentage of significant tests using eMPress with one host per symbiont | P: 5%<br>P+C: 0% | P: 8%<br>P+C: 0% | P: 16%<br>P+C: 0% | P: 18%<br>P+C: 0% | P: 100%<br>P+C: 92% | P: 99%<br>P+C: 34% |
|  | Percentage of significant tests using eMPress with random bifurcations | P: 34%<br>P+C: 0% | P: 23%<br>P+C: 0% | P: 80%<br>P+C: 0% | P: 58%<br>P+C: 0% |  |  |

**Figure 2:**
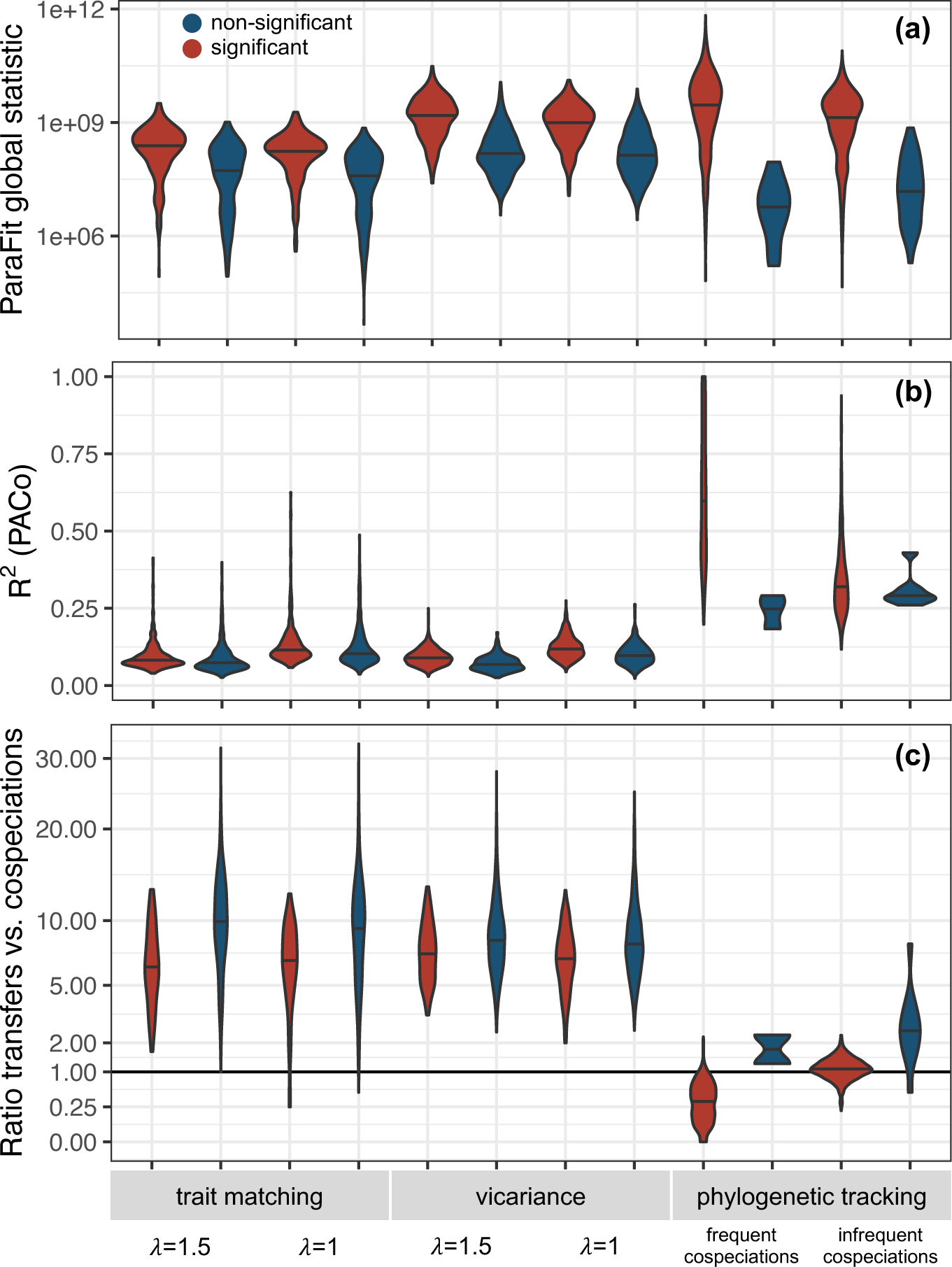
eMPRess correctly distinguishes patterns of phylogenetic congruence (scenario of phylogenetic tracking with frequent cospeciations) from cophylogenetic signal alone (scenarios of trait matching and vicariance), whereas ParaFit cannot and PACo can only in some cases: Distribution of ParaFit global statistics (a), PACo’s R^2^ (b), and the ratio between the number of host transfers and the number of cospeciations in eMPRess reconciliations (c) as a function of the test significance and the simulation scenario of trait matching, vicariance, or phylogenetic tracking. Here, we only reported eMPRess reconciliations obtained when subsampling one host per symbiont species (with relative costs d=4, t=1, and l=1), but similar results were observed when using random bifurcations (Supplementary Fig. S5).

**Figure 3:**
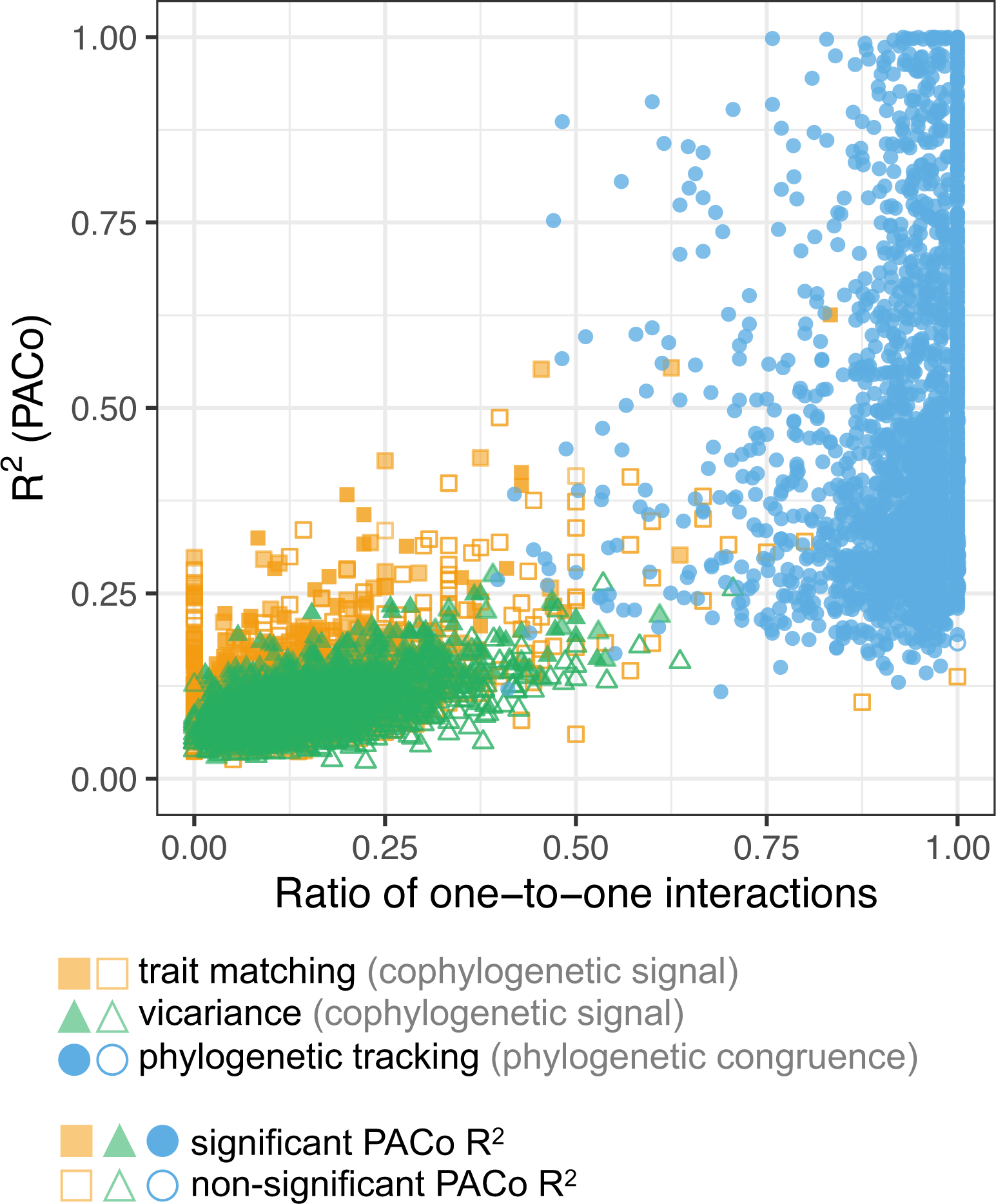
PACo’s statistic (R^2^) tends to increase with the ratio of one-to-one interactions; however, PACo tests tend to be more often significant when the ratio of one-to-one interaction is low under scenarios of trait matching and vicariance.

## Discussion

Using simulations, we have assessed the ability of different cophylogenetic methods to distinguish phylogenetic congruence from cophylogenetic signal. The distinction is important, as these patterns are indicative of different processes, phylogenetic congruence revealing more intricate evolutionary histories between hosts and symbionts than cophylogenetic signal (Fig. 4). We have shown that global-fit methods typically return significant results as soon as there is cophylogenetic signal in species interactions, including in the absence of phylogenetic congruence, meaning that they cannot distinguish the two patterns. In contrast, event-based methods can be used to specifically detect phylogenetic congruence.

**Figure 4:**
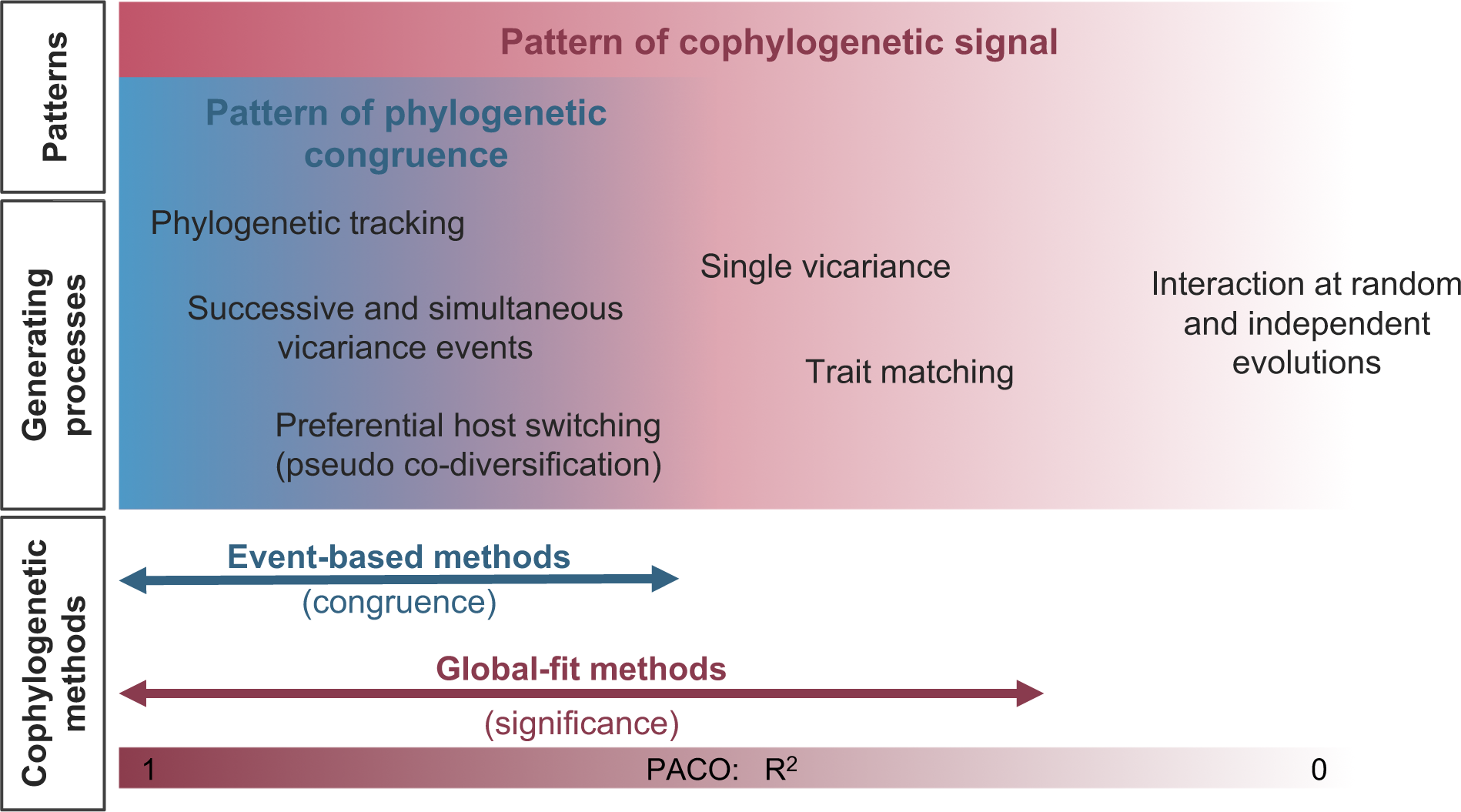
Patterns of cophylogenetic signal and phylogenetic congruence can be generated by various processes: Event-based and global-fit methods differently measure these patterns: Event-based methods can robustly identify phylogenetic congruence, whereas global-fit methods measure cophylogenetic signal. Some of the statistics of global-fit methods (e.g. the R^2^ of PACo) can inform whether the cophylogenetic signal may be due to phylogenetic congruence: a low R^2^ (R^2^<0.25) indicates that there is a cophylogenetic signal but no phylogenetic congruence, whereas a higher R^2^ (R^2^>0.25) suggests there is a cophylogenetic signal and potentially also phylogenetic congruence, but the latter needs to be validated using eMPRess, as R^2^>0.25 also frequently occur in systems without phylogenetic congruence (Fig. 3).

Given that global-fit methods detect cophylogenetic signal rather than phylogenetic congruence, Mantel tests measuring phylogenetic signal in species interactions could be used in place of these methods (Perez-Lamarque et al. 2022b). It would be useful, in the future, to compare the behavior of Mantel tests to global-fit approaches. Regarding event-based methods other than eMPRess, such as TreeMap (Page 1994a, 1995), TreeFitter (Ronquist 2003a), or Jane (Conow et al. 2010), we expect them to have similar behaviors given that they are also cost-based and use maximum parsimony. There are however notable differences between them; for example, eMPRess (like TreeMap or Jane) sets a null cost to cospeciation events, therefore implicitly favoring cospeciation over other reconciliation events (host transfers, intra-host duplication, and symbiont loss), while TreeFitter (Ronquist 2003a) allows setting a positive cost to cospeciation events. Given that cospeciation events may not be that frequent in nature (Ronquist 1995), TreeFitter is probably less likely to overestimate cospeciation events in the reconciliation. More sophisticated probabilistic methods, such as the version of the amalgamated likelihood estimation (ALE) that considers branching orders in addition to tree topology (Szöllősi et al. 2013), may also perform better as it would provide only time-consistent reconciliations (Maestri et al. 2023).

Several recent studies have interpreted significant results of global-fit approaches as evidence for phylogenetic congruence and signs of codiversification (*e.g.* Fuzessy et al. 2022; Suzuki et al. 2022). Our findings suggest that there is evidence for cophylogenetic signal in the studied systems, which is already insightful in itself, but that little can be said about phylogenetic congruence before event-based methods are applied. Applying event-based methods could drastically change the biological conclusions that have been drawn.

Our results suggest that the complementary use of PACo (fast and easy to run) and eMPRess (more computationally intensive but more informative) can be most useful for analyzing cophylogenetic data, with a careful interpretation of the results (Fig. 4). We recommend to begin by using PACo; if the test is not significant, there is neither cophylogenetic signal nor phylogenetic congruence in the data, suggesting independent evolution. If the PACo test is significant but with a low R^2^ (R^2^<0.25), there is a cophylogenetic signal but no congruence, suggesting that evolutionary history played a role in shaping present-day interactions, but that there was not a strong co-dependency during the evolutionary history of the two clades. If the PACo test is significant with a higher R^2^ (R^2^>0.25), there is a cophylogenetic signal, and potentially also phylogenetic congruence, but the latter needs to be validated using eMPRess, as R^2^>0.25 also frequently occur in systems without phylogenetic congruence. Before running eMPRess, if some symbiont species interact with several host species, we recommend randomly sampling one host species per symbiont (rather than, for example, generating random bifurcations in the symbiont tree, which seems to foster false positives). If the eMPRess test supports phylogenetic congruence (significant reconciliation with a larger number of cospeciation events compared with the number of host transfers), this suggests a strong co-dependency during the evolutionary history of the two clades, such as phylogenetic tracking, successive vicariance events, or preferential host switching. As eMPRess uses only the tree topologies and not the branch lengths, no conclusion can be drawn about the concomitance of divergence times in the host and symbionts trees. Additional analyses are thus needed to be able to distinguish phylogenetic congruence with concomitant divergence times (pattern of codiversification arising, *e.g.* from phylogenetic tracking or successive vicariance events) from phylogenetic congruence with non-concomitant divergence times (pattern of pseudo-codiversification arising, *e.g.* from preferential host switching; Ronquist 2003b; de Vienne et al. 2013). One possibility is to check whether eMPRess reconciliations include time-inconsistent host transfers (*i.e.* “back-in-time” transfers between non-contemporary host lineages; Maestri et al. 2023), which would suggest pseudo-codiversification.

An advantage of global-fit methods is that their hypothesis testing is rather flexible. One can easily imagine designing more constrained randomization strategies, for example, to specifically test the influence of biogeography or trait matching on the observed cophylogenetic signal (Perez-Lamarque and Morlon 2023). In parallel, data augmentation and other machine-learning techniques may allow overriding the computational bottleneck that limits the implementation of more complex process-based models. Altogether, such advancements will facilitate linking patterns to processes in cophylogenetic systems.

### Concluding remarks

To conclude, our results imply that using global-fit methods alone is not sufficient to robustly assess a pattern of phylogenetic congruence. In a given cophylogenetic system, finding both significant global-fit tests and significant reconciliations using event-based approaches suggests that there is a pattern of phylogenetic congruence that can be linked to various processes such as phylogenetic tracking, successive vicariance events, or pseudo-codiversification. In contrast, finding significant global-fit tests but no significant reconciliations with event-based approaches suggests that phylogenetic congruence is unlikely; the cophylogenetic signal in this system may rather emerge from processes such as trait matching or biogeographical contingency, but not from phylogenetic tracking or pseudo-codiversification. Clearly distinguishing patterns of cophylogenetic signal and phylogenetic congruence and carefully interpreting outputs of cophylogenetic methods are key if we are to understand the processes that shape present-day communities.

## Acknowledgments

This work was performed using HPC resources from GENCI-IDRIS (Grants 2021-A0100312405 and 2022-AD010313735). The authors acknowledge the Editors Isabel Sanmartín and Lisa Barrow, as well as David Hembry and one anonymous reviewer for insightful comments on an earlier version of the manuscript. We also acknowledge members of the ‘Modeling biodiversity’ lab at IBENS for helpful discussions.

## Supplementary Information

Supplementary information can be found on DRYAD: 10.5061/dryad.7wm37pvzn (temporary link: https://datadryad.org/stash/share/u1-stLek8-xsOgCKnCdGzPXl6KaQopUuY3h1NNOfCeI)

## Code availability

All the codes for generating the simulations and performing the analyses are available through the following link: https://github.com/BPerezLamarque/Scripts/tree/master/Cophylogenetic_signal.

Upon publication, all the codes for generating the simulations and performing the analyses will be available on DRYAD: https://doi.org/10.5061/dryad.7wm37pvzn (temporary link: https://datadryad.org/stash/share/u1-stLek8-xsOgCKnCdGzPXl6KaQopUuY3h1NNOfCeI)

## Author contributions

BPL and HM designed the study, BPL performed the analyses, and both authors wrote the manuscript.

## References

Althoff D.M., Segraves K.A., Johnson M.T.J. 2014. Testing for coevolutionary diversification: Linking pattern with process. Trends Ecol. Evol. 29:82–89.

Balbuena J.A., Míguez-Lozano R., Blasco-Costa I. 2013. PACo: A novel procrustes application to cophylogenetic analysis. PLoS One. 8:e61048.

Bascompte J., Jordano P. 2013. Mutualistic networks. Princeton University Press.

Blasco-Costa I., Hayward A., Poulin R., Balbuena J.A. 2021. Next-generation cophylogeny: unravelling eco-evolutionary processes. Trends Ecol. Evol. 36:907– 918.

Bright M., Bulgheresi S. 2010. A complex journey: transmission of microbial symbionts. Nat Rev Microbiol. 8:218–230.

Conow C., Fielder D., Ovadia Y., Libeskind-Hadas R. 2010. Jane: A new tool for the cophylogeny reconstruction problem. Algorithms Mol. Biol. 5:1–10.

Dismukes W., Braga M.P., Hembry D.H., Heath T.A., Landis M.J. 2022. Cophylogenetic methods to untangle the evolutionary history of ecological interactions. Annu. Rev. Ecol. Evol. Syst. 53:1–24.

Fahrenholz H. 1912. Ectoparasiten und abstammungslehre. Zool. Anz. 41:371–374.

Fuzessy L., Silveira F.A.O., Culot L., Jordano P., Verdú M. 2022. Phylogenetic congruence between Neotropical primates and plants is driven by frugivory. Ecol. Lett. 25:320–329.

Groussin M., Mazel F., Sanders J.G., Smillie C.S., Lavergne S., Thuiller W., Alm E.J. 2017. Unraveling the processes shaping mammalian gut microbiomes over evolutionary time. Nat. Commun. 8:14319.

Harmon L.J., Andreazzi C.S., Débarre F., Drury J., Goldberg E.E., Martins A.B., Melián C.J., Narwani A., Nuismer S.L., Pennell M.W., Rudman S.M., Seehausen O., Silvestro D., Weber M., Matthews B. 2019. Detecting the macroevolutionary signal of species interactions. J. Evol. Biol. 32:769–782.

Hayward A., Poulin R., Nakagawa S. 2021. A broadscale analysis of host-symbiont cophylogeny reveals the drivers of phylogenetic congruence. Ecol. Lett. 24:1681– 1696.

Hembry D.H., Weber M.G. 2020. Ecological interactions and macroevolution: A new field with old roots. Annu. Rev. Ecol. Evol. Syst. 51:215–243.

Hutchinson M.C., Cagua E.F., Balbuena J.A., Stouffer D.B., Poisot T. 2017. paco: implementing Procrustean Approach to Cophylogeny in R. Methods Ecol. Evol. 8:932–940.

Jousselin E., Desdevises Y., Coeur D’Acier A. 2009. Fine-scale cospeciation between Brachycaudus and Buchnera aphidicola: Bacterial genome helps define species and evolutionary relationships in aphids. Proc. R. Soc. B Biol. Sci. 276:187–196.

Legendre P., Desdevises Y., Bazin E. 2002. A statistical test for host-parasite coevolution. Syst. Biol. 51:217–234.

Maestri R., Perez-Lamarque B., Zhukova A., Morlon H. 2023. Recent evolutionary origin and localized diversity hotspots of mammalian coronaviruses. bioRxiv.: 2023.03.09.531875.

Mittelbach G.G., McGill B.J. 2019. Community Ecology. Oxford University Press.

Morand S., Krasnov B.R., Littlewood D.T.J. 2015. Parasite diversity and diversification. Cambridge University Press.

Muchhala N., Thomson J.D. 2009. Going to great lengths: Selection for long corolla tubes in an extremely specialized bat-flower mutualism. Proc. R. Soc. B Biol. Sci. 276:2147–2152.

Page R.D.M. 1994a. Parallel phylogenies: Reconstructing the history of host-parasite assemblages. Cladistics. 10:155–173.

Page R.D.M. 1994b. Maps between trees and cladistic analysis of historical associations among genes, organisms, and areas. Syst. Biol. 43:58–77.

Page R.D.M. 1995. TreeMap. Computer program distributed by the author. University of Glasgow.

Paradis E., Claude J., Strimmer K. 2004. APE: Analyses of phylogenetics and evolution in R language. Bioinformatics. 20:289–290.

Perez-Lamarque B., Krehenwinkel H., Gillespie R.G., Morlon H. 2022a. Limited evidence for microbial transmission in the phylosymbiosis between Hawaiian spiders and their microbiota. mSystems. 7:e01104–21.

Perez-Lamarque B., Maliet O., Pichon B., Selosse M.-A., Martos F., Morlon H. 2022b. Do closely related species interact with similar partners? Testing for phylogenetic signal in bipartite interaction networks. Peer Community J. 2:e59.

Perez-Lamarque B., Morlon H. 2023. Comparing different computational approaches for detecting long-term vertical transmission in host-associated microbiota. Mol. Ecol. 32:6671–6685.

Pichon B., Le Goff R., Morlon H., Perez-Lamarque B. 2023. Telling mutualistic and antagonistic ecological networks apart by learning their multiscale structure. bioRxiv.:2023.04.04.535603.

Poisot T. 2015. When is co-phylogeny evidence of coevolution? Parasite Divers. Diversif. Evol. Ecol. Meets Phylogenetics.:420–433.

Ronquist F. 1995. Reconstructing the history of host-parasite associations using generalised parsimony. Cladistics. 11:73–89.

Ronquist F. 1998. Phylogenetic approaches in coevolution and biogeography. Zool. Scr. 26:313–322.

Ronquist F. 2003a. TreeFitter, version 1.2., Software available from https://sourceforge.net/projects/treefitter/.

Ronquist F. 2003b. Parsimony analysis of coevolving species associations. Tangl. Trees Phylogeny, Cospeciation Coevol.:22–64.

Ronquist F., Nylin S. 1990. Process and pattern in the evolution of species associations. Syst. Zool. 39:323–344.

Sanmartín I., Ronquist F. 2002. New solutions to old problems: widespread taxa, redundant distributions and missing areas in event-based biogeography. Anim. Biodivers. Conserv. 25:75–93.

Sanmartín I., Ronquist F. 2004. Southern hemisphere biogeography inferred by event-based models: Plant versus animal patterns. Syst. Biol. 53:216–243.

Santichaivekin S., Yang Q., Liu J., Mawhorter R., Jiang J., Wesley T., Wu Y.C., Libeskind-Hadas R. 2021. EMPRess: A systematic cophylogeny reconciliation tool. Bioinformatics. 37:2481–2482.

Satler J.D., Herre E.A., Jandér K.C., Eaton D.A.R., Machado C.A., Heath T.A., Nason J.D. 2019. Inferring processes of coevolutionary diversification in a community of Panamanian strangler figs and associated pollinating wasps. Evolution (N. Y). 73:2295–2311.

Su Z.-H., Sasaki A., Kusumi J., Chou P.-A., Tzeng H.-Y., Li H.-Q., Yu H. 2022. Pollinator sharing, copollination, and speciation by host shifting among six closely related dioecious fig species. Commun. Biol. 5:284.

Suzuki T.A., Fitzstevens J.L., Schmidt V.T., Enav H., Huus K.E., Ngwese M.M., Grießhammer A., Pfleiderer A., Adegbite B.R., Zinsou J.F., Esen M., Velavan T.P., Adegnika A.A., Song L.H., Spector T.D., Muehlbauer A.L., Marchi N., Kang H., Maier L., Blekhman R., Ségurel L., Ko G.P., Youngblut N.D., Kremsner P., Ley R.E. 2022. Codiversification of gut microbiota with humans. Science (80-.). 377:1328–1332.

Szöllősi G.J., Rosikiewicz W., Boussau B., Tannier E., Daubin V., Szöllosi G.J., Rosikiewicz W., Boussau B., Tannier E., Daubin V. 2013. Efficient exploration of the space of reconciled gene trees. Syst. Biol. 62:901–912.

de Vienne D.M., Refrégier G., López-Villavicencio M., Tellier A., Hood M.E., Giraud T. 2013. Cospeciation vs host-shift speciation: Methods for testing, evidence from natural associations and relation to coevolution. New Phytol. 198:347–385.

