## Supplementary material for "Distinguishing cophylogenetic signal from phylogenetic congruence clarifies the interplay between evolutionary history and species interactions": Supp Fig

**Supplementary Box (1)**

**Supplementary Methods (1)**

**Supplementary Tables (1-6)**

**Supplementary Figures (1-5)**

**Supplementary References**

**Supplementary Box 1:**

**Host-symbiont bipartite interaction network:** A network representing which symbiont species interact with which host, here encoded by a matrix with host species in rows and symbiont species in columns, and 1/0 representing the presence/absence of interaction, respectively.
**Codiversification/Co-cladogenesis:** Pattern of concomitant diversification events happening in both host and symbiont clades. Codiversification can occur due to processes of phylogenetic tracking or successive vicariance events affecting both clades.
**Coevolution:** Process of reciprocal evolutionary changes induced by selective pressures in two (or more) interacting lineages.
**Cophylogenetic signal:** Pattern depicting the tendency of closely related species to interact with closely related partners.
**Cophylogenetics:** Study of the link between the host and symbiont evolutionary histories and extant interactions.
**Cospeciation:** Concomitant event of host and symbiont speciations.
**Degree of specialization:** Number of species interacting with a given focal species.
**Event-based methods:** Cophylogenetic methods reconciling the host and symbiont phylogenies by fitting reconciliation events (e.g. cospeciation, host transfer, duplication, or loss) on the symbiont phylogeny.
**Global-fit methods:** Cophylogenetic methods evaluating the presence of cophylogenetic signal in a bipartite network.
**Intra-host duplications:** Process of symbiont speciation within a host lineage.
**Host transfer (syn. host switch, host shift):** Process of transmission of a symbiont from a donor host to a receiver host.
**Phylogenetic congruence:** Pattern of high similarity of the phylogenetic trees of interacting host and symbiont clades in terms of topology and relative branch lengths. If host and symbiont divergence times are matching, phylogenetic congruence can correspond to codiversification.
**Phylogenetic signal:** Pattern depicting the tendency of closely related species to have similar traits.
**Phylogenetic tracking:** Process of host speciations generating subsequent symbiont speciations (e.g. because of vertical transmission that isolates symbiont populations between the two daughter host lineages). In addition, symbionts can experience events of host transfers, intra-host duplication, or loss at low frequency. Phylogenetic tracking usually generates a pattern of codiversification.

**Preferential host switching**: Tendency of symbionts to experience host transfers toward closely related host species; when the transfer results in a speciation event in the symbiont lineage, this tends to generate phylogenetic congruence. This does not imply co-diversification though, as the divergence times of the symbionts may be much more recent than those of the hosts. We specifically refer to this resulting pattern as **pseudo-codiversification.**

**Symbiont loss:** Extinction of a symbiont in a given host lineage.

**Tree topology:** Branching pattern of the different nodes of a phylogenetic tree that does not consider branch lengths (*i.e.* evolutionary time).

**Vertical transmission:** Process of symbiont inheritance from host generation to host generation (at the level of individual or at the level of the whole lineage). Over long timescales, it can generate phylogenetic tracking.

**Vicariance:** Formation of biogeographic barriers leading to the isolation of populations and subsequent allopatric speciation. By simultaneously affecting hosts and symbionts, vicariance can lead to cophylogenetic signal, and potentially phylogenetic congruence if events of vicariance occur repetitively.

**Supplementary Methods:**

**Supplementary Methods 1: Simulations of cophylogenetic systems**

All simulations were performed in R (R Core Team 2022).

**Scenario of trait matching:**

In the first simulated scenario, we assumed that host-symbiont interactions at present are more likely between species having complementary traits following a trait-matching expression with a unidimensional continuous trait. We independently simulated two phylogenetic trees for the host and symbiont clades using a birth-death model using the *pbtree* function in the R-package phytools (Revell 2012). To do so, for each simulation, we sampled the number of host and symbiont species uniformly in [10, 50], [51, 100], [101, 150], or [151, 200] to test the effect of clade sizes on the cophylogenetic methods. Given the number of species, we simulated the phylogenetic trees with a speciation rate of 0.1 for the hosts (or 0.3 for the symbionts) and an extinction rate of 0.003 for the hosts. We therefore obtained host and symbiont phylogenetic trees with approximately similar numbers of species, but the ages of the symbiont clades are on average much younger than the host clades (Supplementary Fig. S1).

On each tree, we then independently simulated the evolution of traits modulating present-day interactions using a Brownian motion for trait evolution with the *mvSIM* function in the R-package mvMORPH (Clavel et al. 2015). Brownian motions were simulated using a variance of 1 and an arbitrary ancestral state of 10. As a result, closely-related hosts (resp. symbionts) tend to have similar trait values (*i.e.* phylogenetic signal in species traits). Finally, we assumed that the degree of specialization of the symbionts (*i.e.* the number of hosts that a given symbiont interacts with) followed a Poisson distribution with parameter $\lambda$=1.5. We attributed the host-symbiont interacting pairs following a trait-matching expression by assuming that the probability of an interaction pair is proportional to the inverse of the absolute distance between the host trait ($x_{host\ i}$) and symbiont trait ($x_{symbiont\ j}$):

$$\text{P(interaction between host } i \text{ and symbiont } j) \sim \frac{1}{|x_{host\ i} - x_{symbiont\ j}|}$$

As a result, closely related host species interact with closely related symbiont species (*i.e.* cophylogenetic signal), although the phylogenetic trees are independent (*i.e.* no phylogenetic congruence).

For each range of clade sizes, we performed 250 simulations, generating a total of 1,000 simulations. In addition, we replicated the simulations with fewer host species associated with each symbiont species by using a Poisson distribution with parameter $\lambda$=1 (instead of 1.5).

**Scenario of vicariance:**

In the second simulated scenario, we assumed that host and symbiont species interact at random as long as they occupy the same biogeographic area. At first, all hosts and symbionts simultaneously occupy a single area and diversify independently: we simulated hosts and symbiont clades constituted of 10 to 20 species (using the *pbtree* function with a speciation rate of 0.1 and an extinction rate of 0.003). Then, we simulated an event of vicariance that splits the area into three separate areas and isolates the different host and symbiont species at random. At the vicariance event, we assumed that a given species had (i) a 50% chance of occupying a single area and (ii) a 50% chance of occupying two isolated areas. The latter scenario resulted in allopatric speciations because of the isolation by vicariance. In other words, on average, 50% of the host and symbiont species experienced an event of speciation at the moment of the vicariance event. Following the vicariance, we assumed that dispersion between areas is not possible and each host and symbiont lineage independently diversifies in its area during 5, 10, 15, or 20 Myr, resulting in more or less species-rich host and symbiont clades.

We thus obtained trees presenting a phylogenetic signal in biogeographic repartition (*i.e.* closely related species tend to occupy the same area). Finally, we assumed that the degree of specialization of the symbionts followed a Poisson distribution with parameter $\lambda$=1.5 and randomly attributed host-symbiont interactions within each area: it thus generates cophylogenetic signal. Because both the host and symbiont clades experienced a burst of speciation at the time of vicariance, their diversification dynamic has not been entirely independent. Yet, they diversify independently before and after the vicariance, which avoids phylogenetic congruence between the host and symbiont phylogenies. Therefore, these simulations generate cophylogenetic signal without phylogenetic congruence.

For each range of clade sizes, we performed 250 simulations, generating a total of 1,000 simulations. In addition, we replicated the simulations with fewer host species associated with each symbiont species by using a Poisson distribution with parameter $\lambda$=1 (instead of 1.5).

**Scenario of phylogenetic tracking:**

In the third simulated scenario, we assume that the symbiont diversification tracks the host diversification: symbionts species are vertically transmitted over long-time scales among host lineages and cospeciate at host speciation events resulting in codiversification.

For each simulation, we sampled the number of host species uniformly in [10, 50], [51, 100], [101, 150], or [151, 200] to test the effect of clade sizes on the cophylogenetic methods. Given the number of host species, we simulated the host phylogenetic tree with a speciation rate of 0.1 and an extinction rate of 0.003. Then, we simulated phylogenetic tracking of the symbionts of the host phylogeny using the R-function *sim_microbiota* from the R-package HOME (Perez-Lamarque and Morlon 2019) with a number of host transfers uniformly sampled between 0 and half the number of host species and an intra-host duplication rate of 0.001. When simulating a host transfer from a donor host to a receiving host, we assumed that the symbiont of the receiving host lineage is replaced (Perez-Lamarque and Morlon 2019). In addition, we assumed that host lineages can lose their symbionts at present by simulating symbiont extinctions with a probability of 0.1 in extant host lineages. Under these simulations, we expect phylogenetic congruence between host and symbiont phylogenies, and therefore also cophylogenetic signal.

We replicated these simulations with more host transfers and intra-host duplications relatively to the number of cospeciations: we uniformly sampled the number of host transfers between 50% and 75% of the number of host species and used an intra-host duplication rate of 0.0015. This second set of simulations breaks the phylogenetic congruence as a majority of events correspond to non-cospeciation events.

In all simulations, we considered only binary interactions: a host-symbiont interaction either exists (1) or does not (0). If any, host or symbiont species interacting with no partners were removed from the trees. When comparing simulations, we noticed that the ratio of one-to-one interactions tends to be lower when simulating scenarios of trait matching or vicariance compared with the scenarios of phylogenetic tracking (Supplementary Fig. 2).

**Supplementary Tables:**

**Supplementary Table 1: Global-fit approaches provide similar results when using null model 2:** The table indicates the percentages of simulations that have a significant test of ParaFit or PACo using null model 2 (*i.e.* shuffling at random the host species names). Three scenarios were tested for simulating host-symbiont cophylogenetic systems: (a) present-day interactions dictated by trait matching (with parameter $\lambda$=1.5), (b) present-day interactions at random following a single event of vicariance (with parameter $\lambda$=1.5), or (c) present-day interactions resulting from phylogenetic tracking (with a majority of cospeciation events).

(a) **Simulations of trait matching:**

| **Methods** | **Number of species per clade** | | | |
|---|---|---|---|---|
| | **Between 10 and 50** | **Between 51 and 100** | **Between 101 and 150** | **Between 151 and 200** |
| **Percentage of significant tests using ParaFit** | 15% | 26% | 35% | 40% |
| **Percentage of significant tests using PACo** | 15% | 28% | 36% | 44% |

(b) **Simulations of vicariance:**

| **Methods** | **Time since vicariance (in Myr)** | | | |
|---|---|---|---|---|
| | **5** | **10** | **15** | **20** |
| **Percentage of significant tests using ParaFit** | 11% | 23% | 57% | 83% |
| **Percentage of significant tests using PACo** | 15% | 39% | 72% | 93% |

(c) **Simulations of phylogenetic tracking:**

| **Methods** | **Number of host species** | | | |
|---|---|---|---|---|
| | **Between 10 and 50** | **Between 51 and 100** | **Between 101 and 150** | **Between 151 and 200** |
| **Percentage of significant tests using ParaFit** | 96% | 100% | 100% | 100% |

| | | | | |
|---|---|---|---|---|
| **Percentage of significant tsts using PACo** | 98% | 100% | 100% | 100% |

**Supplementary Table 2: Results obtained with global-fit methods detailed for the different number of species per clade:** The table indicates the percentages of simulations that have a significant test of ParaFit or PACo using null model 1. Three scenarios were tested: (a) present-day interactions dictated by trait matching (with parameter λ=1.5), (b) present-day interactions at random following a single event of vicariance (with parameter λ=1.5), or (c) present-day interactions resulting from phylogenetic tracking (with a majority of cospeciation events).

(a) **Simulations of trait matching:**

| **Methods** | **Number of species per clade** | | | |
|---|---|---|---|---|
| | **Between 10 and 50** | **Between 51 and 100** | **Between 101 and 150** | **Between 151 and 200** |
| **Percentage of significant tests using ParaFit** | 23% | 34% | 48% | 51% |
| **Percentage of significant tests using PACo** | 31% | 44% | 60% | 66% |

(b) **Simulations of vicariance:**

| **Methods** | **Time since vicariance (in Myr)** | | | |
|---|---|---|---|---|
| | **5** | **10** | **15** | **20** |
| **Percentage of significant tests using ParaFit** | 15% | 34% | 71% | 90% |
| **Percentage of significant tests using PACo** | 25% | 60% | 88% | 98% |

(c) **Simulations of phylogenetic tracking:**

| **Methods** | **Number of host species** | | | |
|---|---|---|---|---|
| | **Between 10 and 50** | **Between 51 and 100** | **Between 101 and 150** | **Between 151 and 200** |
| **Percentage of significant tests using ParaFit** | 96% | 100% | 100% | 100% |
| **Percentage of significant tests using PACo** | 98% | 100% | 100% | 100% |

**Supplementary Table 3: Global-fit approaches tend to be less significant when simulating cophylogenetic signal with symbiont species interacting on average with fewer hosts (more one-to-one interactions), while eMPRess provides qualitatively similar results.** The following tables indicate the percentages of simulations that have a significant test of ParaFit, PACo, or eMPRess when simulating symbiont species with fewer associated host species (i.e. using a Poisson distribution with a parameter $\lambda$=1 instead of $\lambda$=1.5 for the number of associated hosts per symbiont species). Two scenarios were tested: (a) present-day interactions dictated by trait matching and (b) present-day interactions at random following a single event of vicariance.
For the global-fit approaches, the results were obtained with null model 1 (results are qualitatively similar for null model 2; not shown). For eMPRess, host-symbiont reconciliations were run with the following relative costs: d=4, t=1, and l=1 for duplications, host transfers, and losses, respectively. We reported the percentage of significant reconciliations based on permutations alone (P) and based on permutations presenting more cospeciation events than host transfer events (P+C). We consider eMPRess to support phylogenetic congruence when the conditions P and C are met.

**(a) Simulations of trait matching:**

| Methods | Number of species per clade | | | |
|---|---|---|---|---|
| | **Between 10 and 50** | **Between 51 and 100** | **Between 101 and 150** | **Between 151 and 200** |
| **Percentage of significant tests using ParaFit** | 16% | 24% | 31% | 44% |
| **Percentage of significant tests using PACo** | 24% | 41% | 50% | 66% |
| **Percentage of significant tests using eMPRess with one host per symbiont** | P: 6%<br>P+C: 1% | P: 7%<br>P+C: 0% | P: 10%<br>P+C: 0% | P: 11%<br>P+C: 0% |
| **Percentage of significant tests using eMPRess with random bifurcations** | P: 18%<br>P+C: 0% | P: 18%<br>P+C: 0% | P: 24%<br>P+C: 0% | P: 32%<br>P+C: 0% |

**(b) Simulations of vicariance:**

| Methods | Time since vicariance (in Myr) | | | |
|---|---|---|---|---|
| | **5** | **10** | **15** | **20** |
| **Percentage of significant tests using ParaFit** | 9% | 21% | 46% | 75% |
| **Percentage of significant tests using PACo** | 18% | 42% | 80% | 95% |
| **Percentage of significant tests using eMPRess with one host per symbiont** | P: 8%<br>P+C: 0% | P: 15%<br>P+C: 0% | P: 19%<br>P+C: 0% | P: 32%<br>P+C: 0% |
| **Percentage of significant tests using eMPRess with random bifurcations** | P: 35%<br>P+C: 0% | P: 51%<br>P+C: 0% | P: 69%<br>P+C: 0% | P: 76%<br>P+C: 0% |

**Supplementary Table 4: On trait matching simulations, eMPRess gave qualitatively similar results for different cost values for duplications (d), host transfers (t), and losses (l).** These tables indicate the percentages of simulations that present a significant host-symbiont reconciliation using eMPRess when simulating present-day interactions dictated by trait matching (λ=1.5) with different numbers of host and symbiont species per clade. We reported the percentage of significant reconciliations based on permutations alone (P) and based on permutations presenting more cospeciation events than host transfer events (P+C). We consider eMPRess to support phylogenetic congruence when conditions P and C are met.

**(a) Percentage of significant tests using eMPRess with one host per symbiont**

| eMPRess cost values | Number of species per clade | | | |
|---|---|---|---|---|
| | **[10 ; 50]** | **[51 ; 100]** | **[101 ; 150]** | **[151 ; 200]** |
| **d=1, t=1, l=1** | P: 3%<br>P+C: 0% | P: 8%<br>P+C: 0% | P: 7%<br>P+C: 0% | P: 13%<br>P+C: 0% |
| **d=4, t=1, l=1** | P: 6%<br>P+C: 0% | P: 6%<br>P+C: 0% | P: 4%<br>P+C: 0% | P: 5%<br>P+C: 0% |
| **d=2, t=1, l=2** | P: 5%<br>P+C: 0% | P: 7%<br>P+C: 0% | P: 5%<br>P+C: 0% | P: 7%<br>P+C: 0% |
| **d=4, t=2, l=1** | P: 8%<br>P+C: 0% | P: 11%<br>P+C: 0% | P: 8%<br>P+C: 0% | P: 13%<br>P+C: 0% |
| **d=2, t=3, l=1** | P: 5%<br>P+C: 1% | P: 10%<br>P+C: 0% | P: 13%<br>P+C: 0% | P: 20%<br>P+C: 0% |

**(b) Percentage of significant tests using eMPRess with random bifurcations**

| eMPRess cost values | Number of species per clade | | | |
|---|---|---|---|---|
| | **[10 ; 50]** | **[51 ; 100]** | **[101 ; 150]** | **[151 ; 200]** |
| **d=1, t=1, l=1** | P: 12%<br>P+C: 0% | P: 18%<br>P+C: 0% | P: 26%<br>P+C: 0% | P: 30%<br>P+C: 0% |
| **d=4, t=1, l=1** | P: 24%<br>P+C: 0% | P: 31%<br>P+C: 0% | P: 37%<br>P+C: 0% | P: 42%<br>P+C: 0% |
| **d=2, t=1, l=2** | P: 18%<br>P+C: 0% | P: 24%<br>P+C: 0% | P: 33%<br>P+C: 0% | P: 37%<br>P+C: 0% |
| **d=4, t=2, l=1** | P: 26%<br>P+C: 0% | P: 34%<br>P+C: 0% | P: 42%<br>P+C: 0% | P: 49%<br>P+C: 0% |
| **d=2, t=3, l=1** | P: 11%<br>P+C: 0% | P: 23%<br>P+C: 0% | P: 34%<br>P+C: 0% | P: 42%<br>P+C: 0% |

**Supplementary Table 5: On vicariance simulations, eMPRess gave qualitatively similar results for different cost values for duplications (d), host transfers (t), and losses (l), with the exception of high costs for transfers (t).** These tables indicate the percentages of simulations that present a significant host-symbiont reconciliation using eMPRess when simulating present-day interactions at random following a single event of vicariance ($\lambda$=1.5) with different times since the event of vicariance. We reported the percentage of significant reconciliations based on permutations alone (P) and based on permutations presenting more cospeciation events than host transfer events (P+C). We consider eMPRess to support phylogenetic congruence when conditions P and C are met.

**(a) Percentage of significant tests using eMPRess with one host per symbiont**

| **eMPRess cost values** | **Time since vicariance (in Myr)** | | | |
|---|---|---|---|---|
| | **5** | **10** | **15** | **20** |
| **d=1, t=1, l=1** | P: 18%<br>P+C: 0% | P: 25%<br>P+C: 0% | P: 36%<br>P+C: 0% | P: 56%<br>P+C: 0% |
| **d=4, t=1, l=1** | P: 8%<br>P+C: 0% | P: 14%<br>P+C: 0% | P: 18%<br>P+C: 0% | P: 26%<br>P+C: 0% |
| **d=2, t=1, l=2** | P: 11%<br>P+C: 0% | P: 20%<br>P+C: 0% | P: 24%<br>P+C: 0% | P: 36%<br>P+C: 0% |
| **d=4, t=2, l=1** | P: 17%<br>P+C: 0% | P: 30%<br>P+C: 0% | P: 47%<br>P+C: 0% | P: 64%<br>P+C: 0% |
| **d=2, t=3, l=1** | P: 24%<br>P+C: 0% | P: 45%<br>P+C: 0% | P: 74%<br>P+C: 0% | P: 89%<br>P+C: 0% |

**(b) Percentage of significant tests using eMPRess with random bifurcations**

| **eMPRess cost values** | **Time since vicariance (in Myr)** | | | |
|---|---|---|---|---|
| | **5** | **10** | **15** | **20** |
| **d=1, t=1, l=1** | P: 56%<br>P+C: 0% | P: 72%<br>P+C: 0% | P: 83%<br>P+C: 0% | P: 88%<br>P+C: 0% |
| **d=4, t=1, l=1** | P: 64%<br>P+C: 0% | P: 80%<br>P+C: 0% | P: 83%<br>P+C: 0% | P: 92%<br>P+C: 0% |
| **d=2, t=1, l=2** | P: 62%<br>P+C: 0% | P: 75%<br>P+C: 0% | P: 83%<br>P+C: 0% | P: 91%<br>P+C: 0% |
| **d=4, t=2, l=1** | P: 78%<br>P+C: 0% | P: 90%<br>P+C: 0% | P: 98%<br>P+C: 0% | P: 99%<br>P+C: 0% |
| **d=2, t=3, l=1** | P: 66%<br>P+C: 0% | P: 90%<br>P+C: 0% | P: 98%<br>P+C: 0% | P: 99%<br>P+C: 0% |

**Supplementary Table 6: On phylogenetic tracking simulations with a majority of cospeciation events, eMPRess gave qualitatively similar results for different cost values for duplications (d), host transfers (t), and losses (l). This** table indicates the percentages of simulations that present a significant host-symbiont reconciliation using eMPRess when simulating present-day interactions resulting from phylogenetic tracking with different numbers of host species per clade. We reported the percentage of significant reconciliations based on permutations alone (P) and based on permutations presenting more cospeciation events than host transfer events (P+C). We consider eMPRess to support phylogenetic congruence when conditions P and C are met.

| **eMPRess cost values** | **Number of host species** | | | |
|---|---|---|---|---|
| | **Between 10 and 50** | **Between 51 and 100** | **Between 101 and 150** | **Between 151 and 200** |
| **d=1, t=1, l=1** | P: 99%<br>P+C: 82% | P: 100%<br>P+C: 92% | P: 100%<br>P+C: 93% | P: 100%<br>P+C: 94% |
| **d=4, t=1, l=1** | P: 99%<br>P+C: 84% | P: 100%<br>P+C: 92% | P: 100%<br>P+C: 94% | P: 100%<br>P+C: 96% |
| **d=2, t=1, l=2** | P: 99%<br>P+C: 76% | P: 100%<br>P+C: 82% | P: 100%<br>P+C: 82% | P: 100%<br>P+C: 82% |
| **d=4, t=2, l=1** | P: 100%<br>P+C: 88% | P: 100%<br>P+C: 96% | P: 100%<br>P+C: 99% | P: 100%<br>P+C: 100% |
| **d=2, t=3, l=1** | P: 100%<br>P+C: 87% | P: 100%<br>P+C: 95% | P: 100%<br>P+C: 97% | P: 100%<br>P+C: 98% |

**Supplementary Table 7: When simulating phylogenetic tracking with less cospeciation events, eMPRess tests are significant but not congruent (estimated numbers of transfers are larger than number of cospeciation events).**

The following table indicates the percentages of simulations that have a significant test of ParaFit, PACo, or eMPRess when simulating phylogenetic tracking with less cospeciation events (and more host transfers and intra-host duplications that dampen the phylogenetic congruence).

For the global-fit approaches, the results were obtained with null model 1 (results are qualitatively similar for null model 2; not shown). For eMPRess, host-symbiont reconciliations were run with the following relative costs: d=4, t=1, and l=1 for duplications, host transfers, and losses, respectively. We reported the percentage of significant reconciliations based on permutations alone (P) and based on permutations presenting more cospeciation events than host transfer events (P+C). We consider eMPRess to support phylogenetic congruence when the conditions P and C are met.

| **Methods** | **Number of host species per clade** | | | |
|---|---|---|---|---|
| | **Between 10 and 50** | **Between 51 and 100** | **Between 101 and 150** | **Between 151 and 200** |
| **Percentage of significant tests using ParaFit** | 86% | 98% | 100% | 100% |
| **Percentage of significant tests using PACo** | 97% | 100% | 100% | 100% |
| **Percentage of significant tests using eMPRess** | P: 95%<br>P+C: 52% | P: 100%<br>P+C: 41% | P: 100%<br>P+C: 25% | P: 100%<br>P+C: 19% |

**Supplementary Figures:**

**Supplementary Figure 1:** Characteristics of the three types of simulations:
(a) Histogram of the number of host species interacting with at least one symbiont.
(b) Histogram of the number of symbiont species interacting with at least one host.
(c) Histogram of the host clade ages.
(d) Histogram of symbiont clade ages.
Three simulated scenarios are represented: (a) present-day interactions dictated by trait matching (with parameter λ=1.5), (b) present-day interactions at random following a single vicariance event (with parameter λ=1.5), or (c) present-day interactions resulting from phylogenetic tracking (with a majority of cospeciation events).

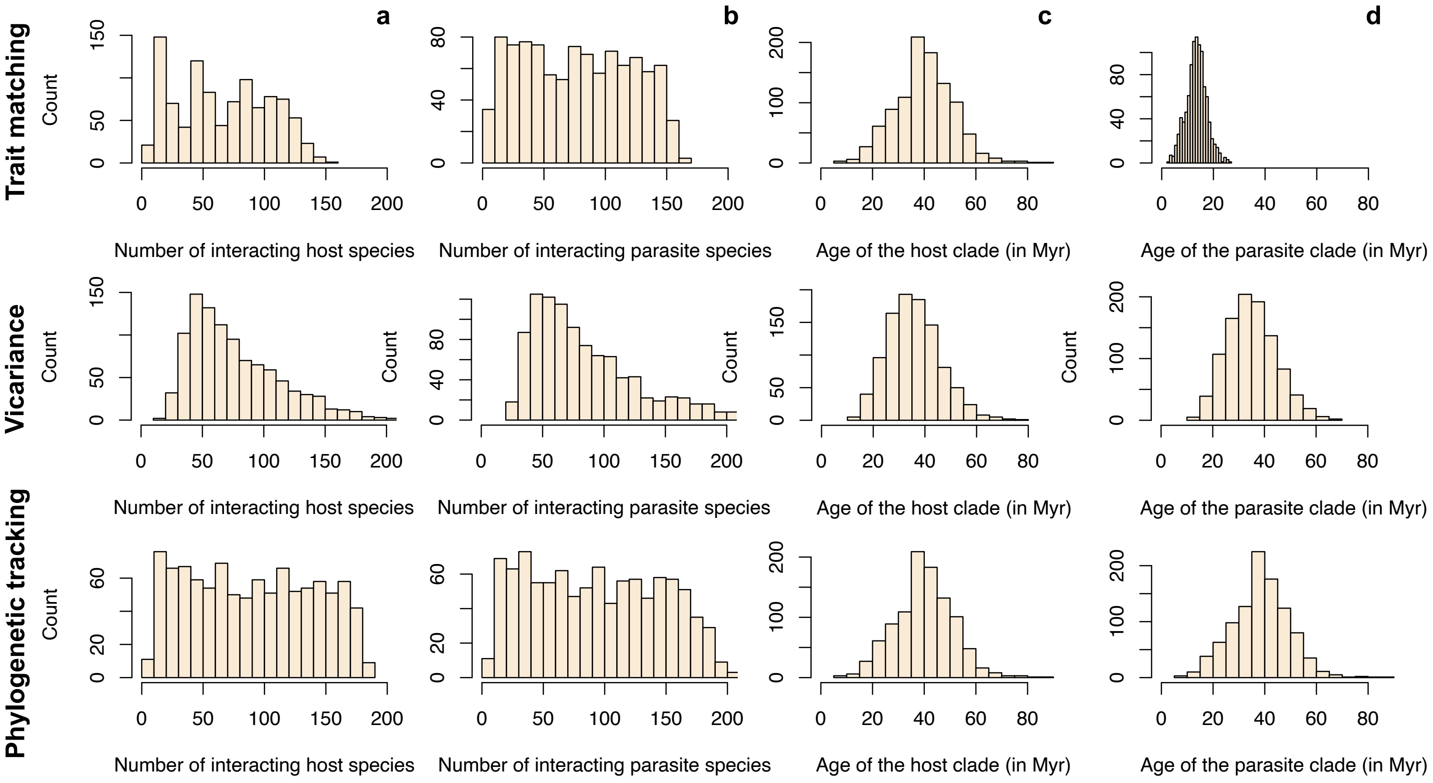

**Supplementary Figure 2: The ratio of one-to-one interactions tends to be lower when simulating trait matching or vicariance.**

Boxplots present the median surrounded by the first and third quartiles, and whiskers extend to the extreme values but no further than 1.5 of the interquartile range.

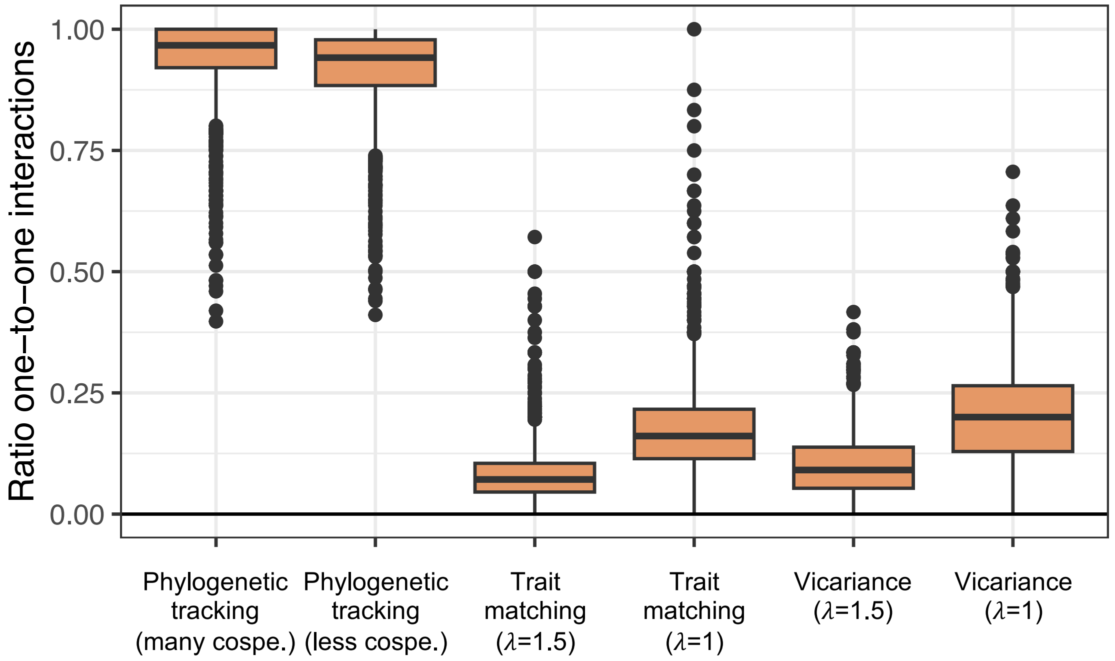

**Supplementary Figure 3: The ParaFit global statistic tends to increase with the total number of species (a) and conversely, it tends to decrease and become non-significant with the ratio of one-to-one interactions (b & c; generalized linear models (GLM): p-values<0.05).** The significance of each ParaFit test was evaluated using null model 1.

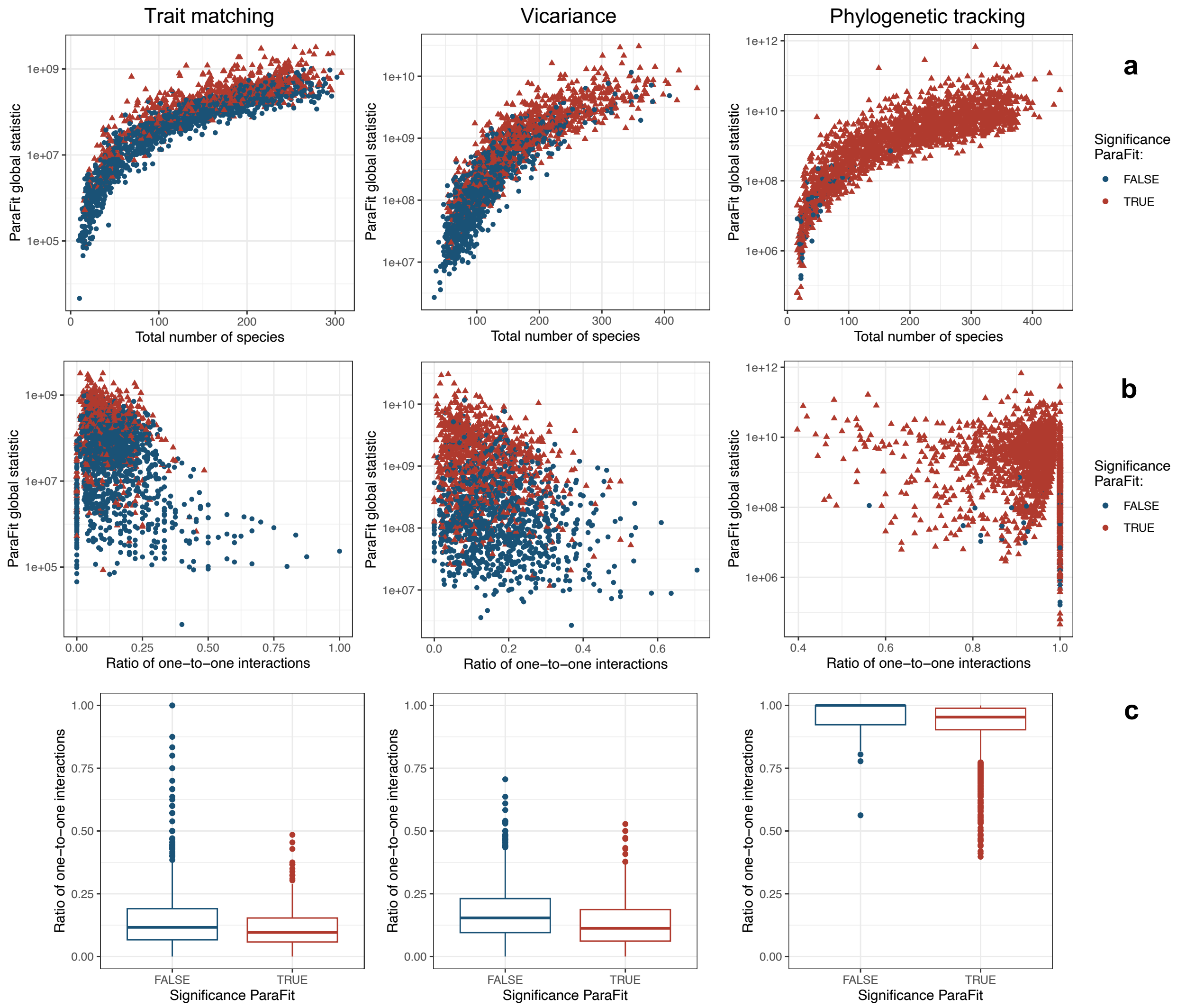

**Supplementary Figure 4: The PACo statistic ($R^2$) does not vary with the total number of species (a) and is generally higher in the presence of phylogenetic congruence (*i.e.* in simulated scenarios with phylogenetic tracking). In simulated scenarios of trait matching and vicariance, it tends to become non-significant for high ratios of one-to-one interactions (b & c; generalized linear models (GLM): p-values<0.05).** The significance of each PACo test was evaluated using null model 1.

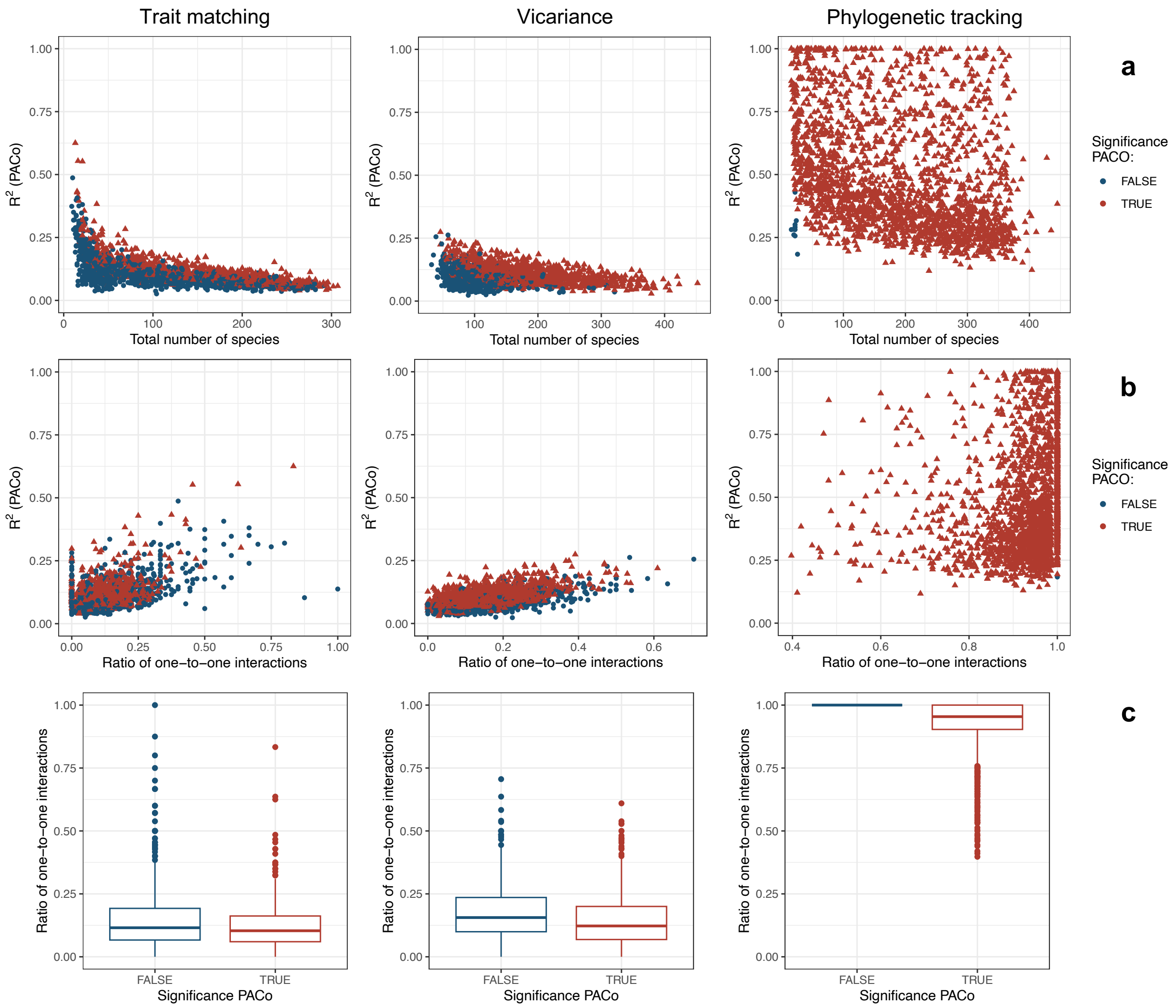

**Supplementary Figure 5: The number of host transfers is much higher than the number of cospeciations for “trait matching” and “vicariance” simulations, while it tends to be lower for “phylogenetic tracking” simulations.**

Boxplots present the median surrounded by the first and third quartiles, and whiskers extend to the extreme values but no further than 1.5 of the interquartile range. Each grey line corresponds to one simulation.

**(a) Percentage of significant tests using eMPRess with one host per symbiont**

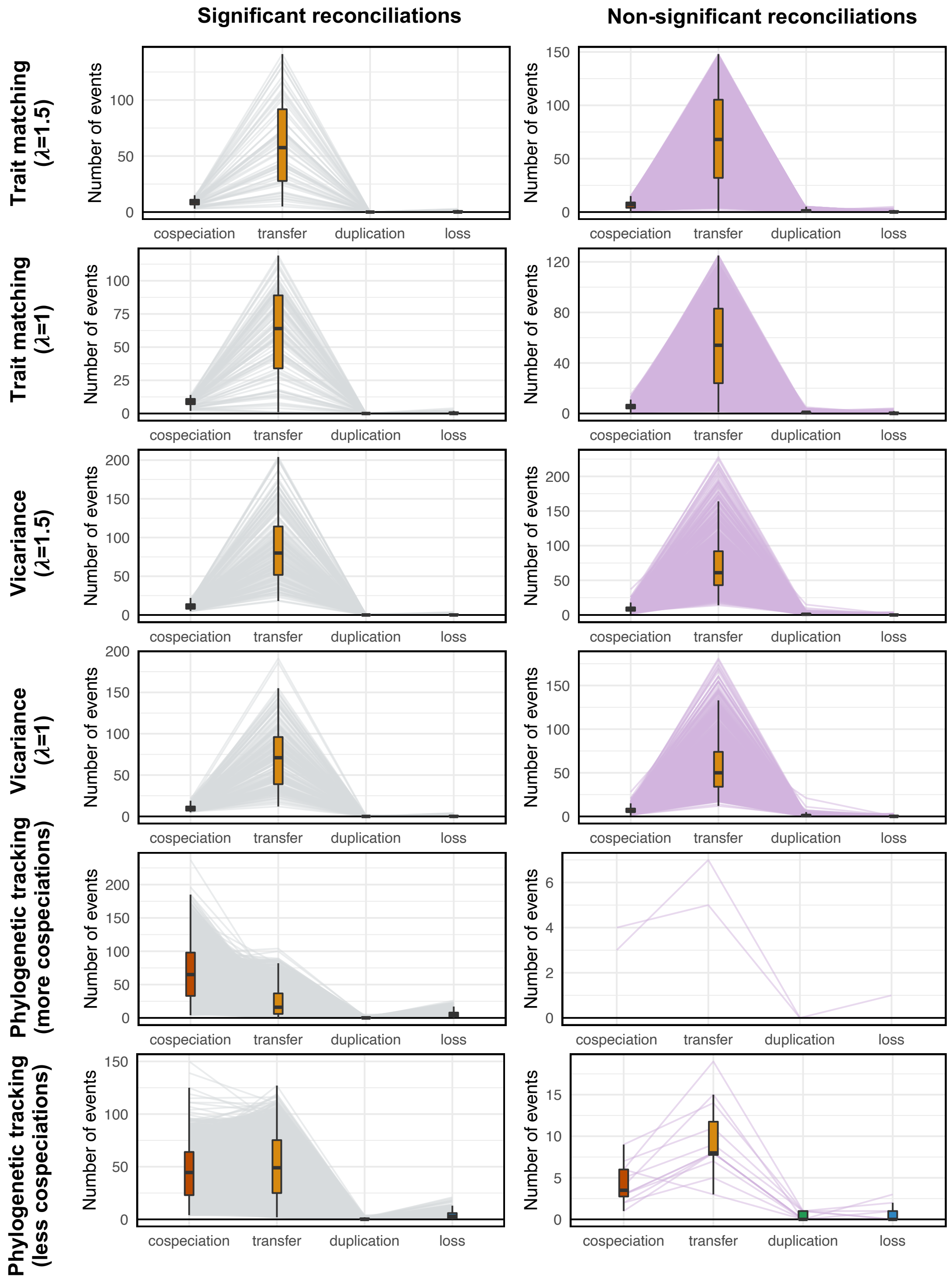

**(b) Percentage of significant tests using eMPRess with random bifurcations**

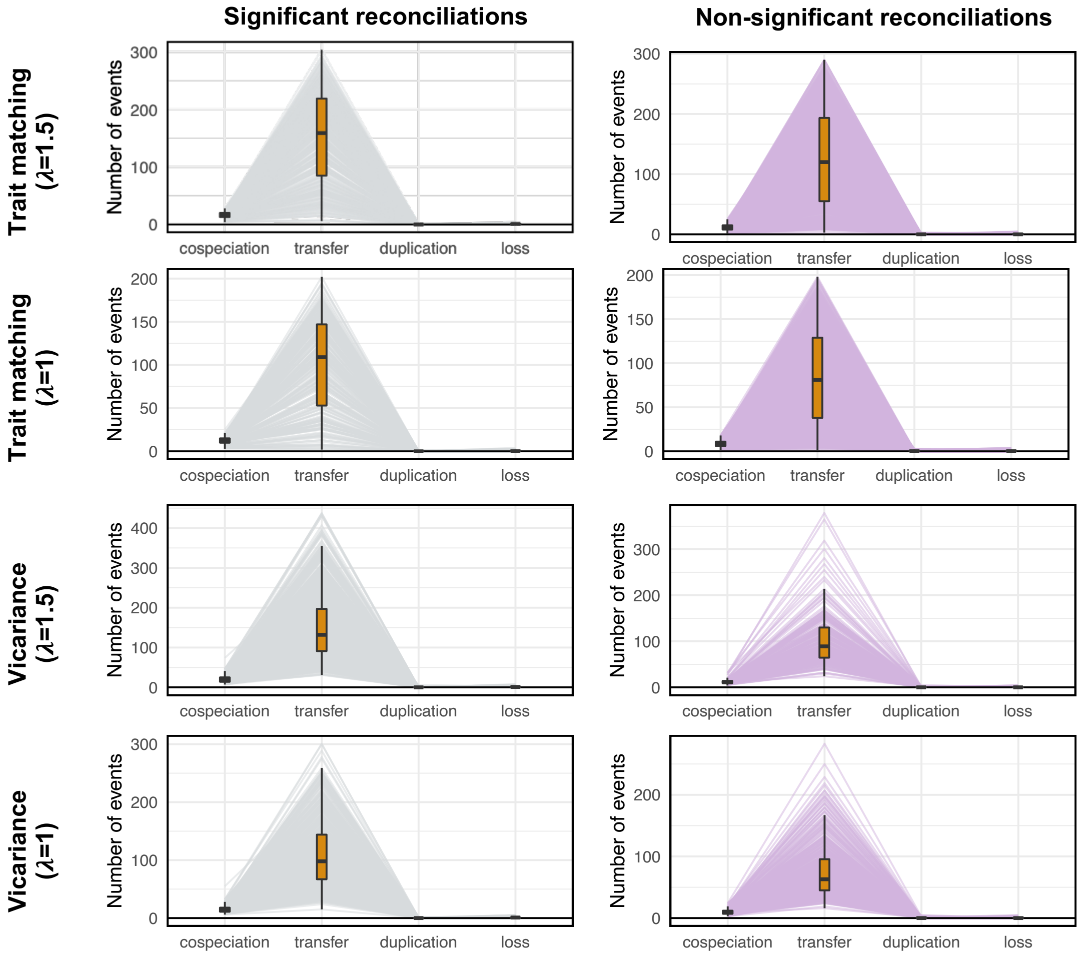

**Supplementary references:**


Clavel J., Escarguel G., Merceron G. 2015. mvMORPH: An R package for fitting multivariate evolutionary models to morphometric data. Methods Ecol. Evol. 6:1311–1319.

Perez-Lamarque B., Morlon H. 2019. Characterizing symbiont inheritance during host–microbiota evolution: Application to the great apes gut microbiota. Mol. Ecol. Resour. 19:1659–1671.

R Core Team. 2022. R: A language and environment for statistical computing. .

Revell L.J. 2012. phytools: An R package for phylogenetic comparative biology (and other things). Methods Ecol. Evol. 3:217–223.
